# Cryo-EM phase-plate images reveal unexpected levels of apparent specimen damage

**DOI:** 10.1101/2024.08.04.606536

**Authors:** Jonathan Remis, Petar N. Petrov, Jessie T Zhang, Jeremy J. Axelrod, Hang Cheng, Shahar Sandhaus, Holger Mueller, Robert M. Glaeser

## Abstract

Apoferritin (apoF) is commonly used as a test specimen in single-particle electron cryo-microscopy (cryo-EM), since it consistently produces density maps that go to 3 Å resolution or higher. When we imaged apoF with a laser phase plate (LPP), however, we observed more severe particle-to-particle variation in the images than we had previously thought to exist. Similarly, we found that images of ribulose bisphosphate carboxylase/oxygenase (rubisco) also exhibited a much greater amount of heterogeneity than expected. By comparison to simulations of images, we verified that the heterogeneity is not explained by the known features of the LPP, shot noise, or differences in particle orientation. We also demonstrate that our specimens are comparable to those previously used in the literature, based on using the final-reconstruction resolution as the metric for evaluation. All of this leads us to the hypothesis that the heterogeneity is due to damage that has occurred either during purification of the specimen or during preparation of the grids. It is not, however, our goal to explain the causes of heterogeneity; rather, we report that using the LPP has made the apparent damage too obvious to be ignored. In hindsight, similar heterogeneity can be seen in images of apoF and the 20S proteasome which others had recorded with a Volta phase plate. We therefore conclude that the increased contrast of phase-plate images (at low spatial frequencies) should also make it possible to visualize, on a single-particle basis, various forms of biologically functional heterogeneity in structure that had previously gone unnoticed.

**GRAPHICAL ABSTRACT:** 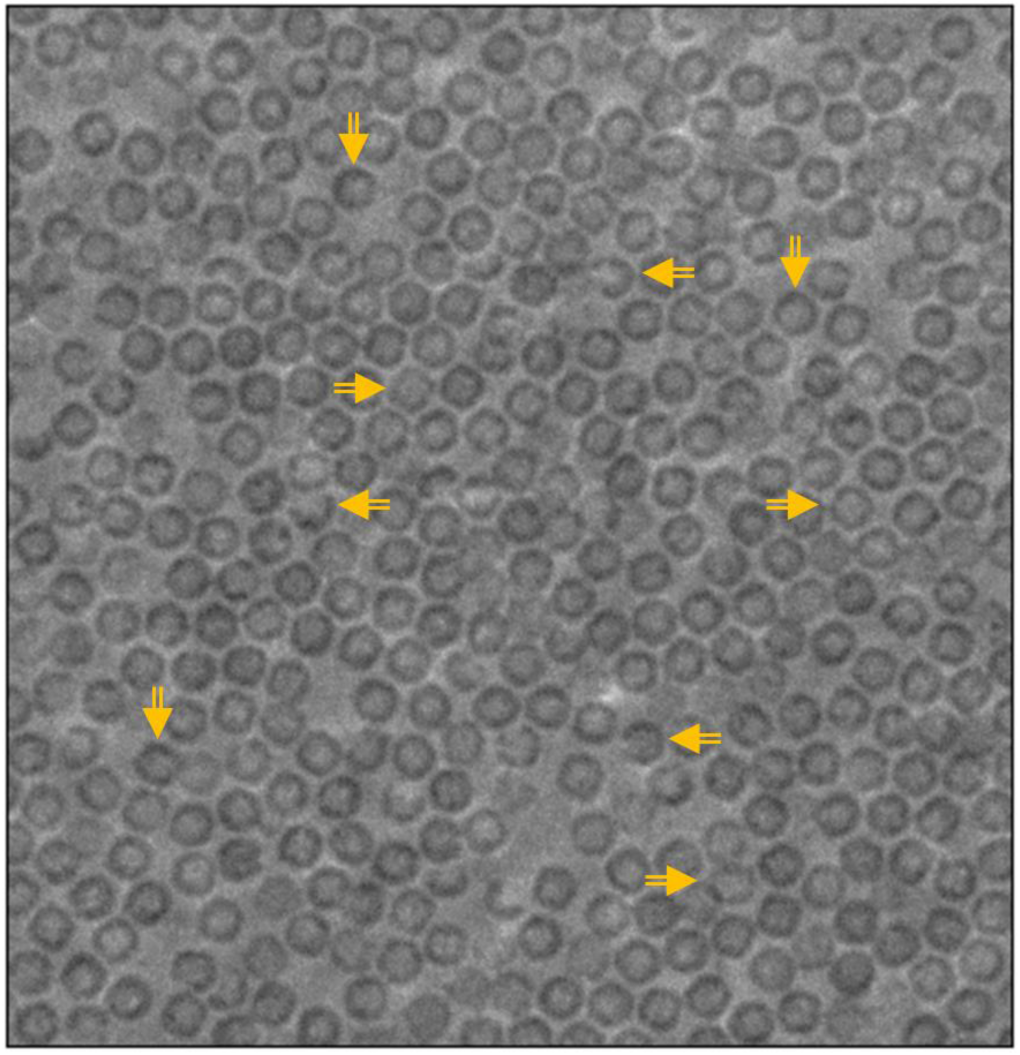

**HIGHLIGHTS:**

- Phase plates recover low-frequency information with significantly improved SNR
- Laser phase-plate images reveal unexpected amounts of structural heterogeneity
- In retrospect, similar heterogeneity can also be seen in Volta phase-plate images
- Particle heterogeneity produces “structural noise”, which may diminish map quality

## INTRODUCTION

Phase plates (Axelrod et al., 2024; Glaeser, 2013) significantly increase the amount of signal that is recovered in single-particle electron cryo-microscopy (cryo-EM) (Glaeser et al., 2021) at low spatial frequencies. The laser phase plate (Schwartz et al., 2019) offers stable and reproducible imaging properties. It employs a Fabry-Perot cavity to build up about 75 kW of circulating power in an optical standing wave, while – at the same time – focusing the light at the center of the electron diffraction pattern (Axelrod et al., 2023; Turnbaugh et al., 2021). The resulting interaction between electrons and light then produces a desirable phase shift between the unscattered and elastically-scattered portions of the diffracted-electron wave function. Use of the LPP is thus expected to considerably improve the ability to visualize structural heterogeneity.

Here, we note that images recorded with the LPP make it possible to recognize what appears to be much more frequent and highly variable instances of particle damage than what is generally thought to exist in the types of specimens we used. When using the LPP to image apoferritin (apoF), which is commonly employed as a high-resolution test specimen (Nakane et al., 2020; Yip et al., 2020; Zhang et al., 2020), we found significant amounts of apparent damage to individual particles. For example, the projected perimeters of individual particles appear to be broken or incomplete, while many other particles show non-uniform or excess density around their perimeters. We verified that these specimens are not atypical – by the conventional, resolution-based standard of the field – since they produce refined 3-D maps at high resolution when imaged both with and without the LPP.

We also observed comparable heterogeneity when imaging ribulose-1,5-bisphosphate carboxylase (rubisco) particles with the LPP. 2-D projections of many particles, even many that contributed to the high-resolution reconstruction, had regions of higher or lower mass density than expected for intact proteins in a thin layer of vitreous ice. As with apoF, our rubisco specimens produced a high-resolution 3-D map when imaged both with and without the LPP, indicating again that the quality of our specimens is consistent with the standards of the field.

In both cases, a comparison of the experimental LPP images to simulations shows that the observed heterogeneity exceeds that expected for undamaged particles. Our simulations took into account effects attributable to the LPP (including phase shifts produced by the optical standing wave (Schwartz et al., 2019) and an additional contribution to the CTF envelope function (Axelrod et al., 2023)), shot noise, and variability in the particle orientation. The discrepancy between simulation and experiment leads us to hypothesize that the apparent heterogeneity reflects some previously unexpected structural features of the specimen, such as damaged protein particles (e.g., at the air-water interface), the presence of protein fragments, or unevenness of the ice.

Such heterogeneity may have been overlooked in conventional cryo-EM images where lower contrast and SNR at low spatial frequencies make it more difficult to distinguish between undamaged and moderately damaged particles. Furthermore, moderate damage might be averaged out during 3-D classification of highly symmetric particles if, for example, only part of a particle is affected. Indeed, inspecting the particle images that were included in the final sets used to produce high-resolution density maps revealed that many of them exhibit structural heterogeneity that is not present in simulated images of undamaged particles.

In retrospect, similar levels of heterogeneity can be seen in images recorded previously with a Volta phase plate, both for apoferritin (Fan et al., 2017; Li et al., 2019) and for the 20S proteasome (Danev et al., 2017). We thus conclude that the increased capability to detect heterogeneity in the specimen (including damage) results from the increase in signal at low spatial frequencies that phase plates provide.

In addition, we report that LPP images of vitrified apoF and rubisco both exhibit a previously unrecognized, low-frequency variation (mottling) of the background of cryo-EM images. Particle heterogeneity and background-mottling both represent unwanted structural noise, which can have an adverse effect on translational and angular alignment of particles, as has been pointed out by (Baxter et al., 2009). We thus suggest that developing methods of sample preparation that, if possible, eliminate these previously-unrecognized defects is likely to bring single-particle cryo-EM even closer to the theoretical limits estimated in the section headed “Averaging particle images” of (Rosenthal and Henderson, 2003).

## MATERIALS AND METHODS

### Cryo-EM sample preparation

VitroEase^TM^ Apoferritin Standard (ThermoFisher Scientific) was used, as supplied, at a concentration of 3.5-4.0 mg/mL. *Halothiobacillus neapolitanos* recombinant rubisco was expressed in *E. coli* BL21-AI cells and purified as described previously by (Oltrogge et al., 2020). Aliquots of rubisco were diluted to 1.3 mg/mL before use.

Samples were vitrified in a Mark IV Vitrobot (Thermo Fisher Scientific), using a chamber temperature of 4 °C and a relative humidity setting of 100 %. The sample was applied in aliquots of 3 µL to Quantifoil R 1.2/1.3 holey carbon (copper) grids (Quantifoil microtools), which were rendered hydrophilic before use by glow-discharge treatment (Pelco easiGlow). Following incubation for 15 s after depositing samples, the grids were blotted for 3 s and then plunged into liquid ethane.

### Data collection

Movies of vitrified samples (Campbell et al., 2012) were recorded using automated, low-dose imaging conditions. External python scripting control of the microscope within SerialEM (Mastronarde, 2005) was used to ensure, before recording each image, that the unscattered electron beam remained properly centered on an antinode of the LPP.

As summarized in **Table 1**, movies were recorded at a nominal (microscope) magnification of 49,000 on a K2 camera (which contributed a calibrated post-magnification factor) operating in counting mode with an effective pixel size of 0.7 Å at the specimen. The total dose was either 37.5 e/Å^2^ or 50.0 e/Å^2^ with a dose per frame of either 0.25 or 1.0 e/Å^2^, respectively.

**Table 1.**
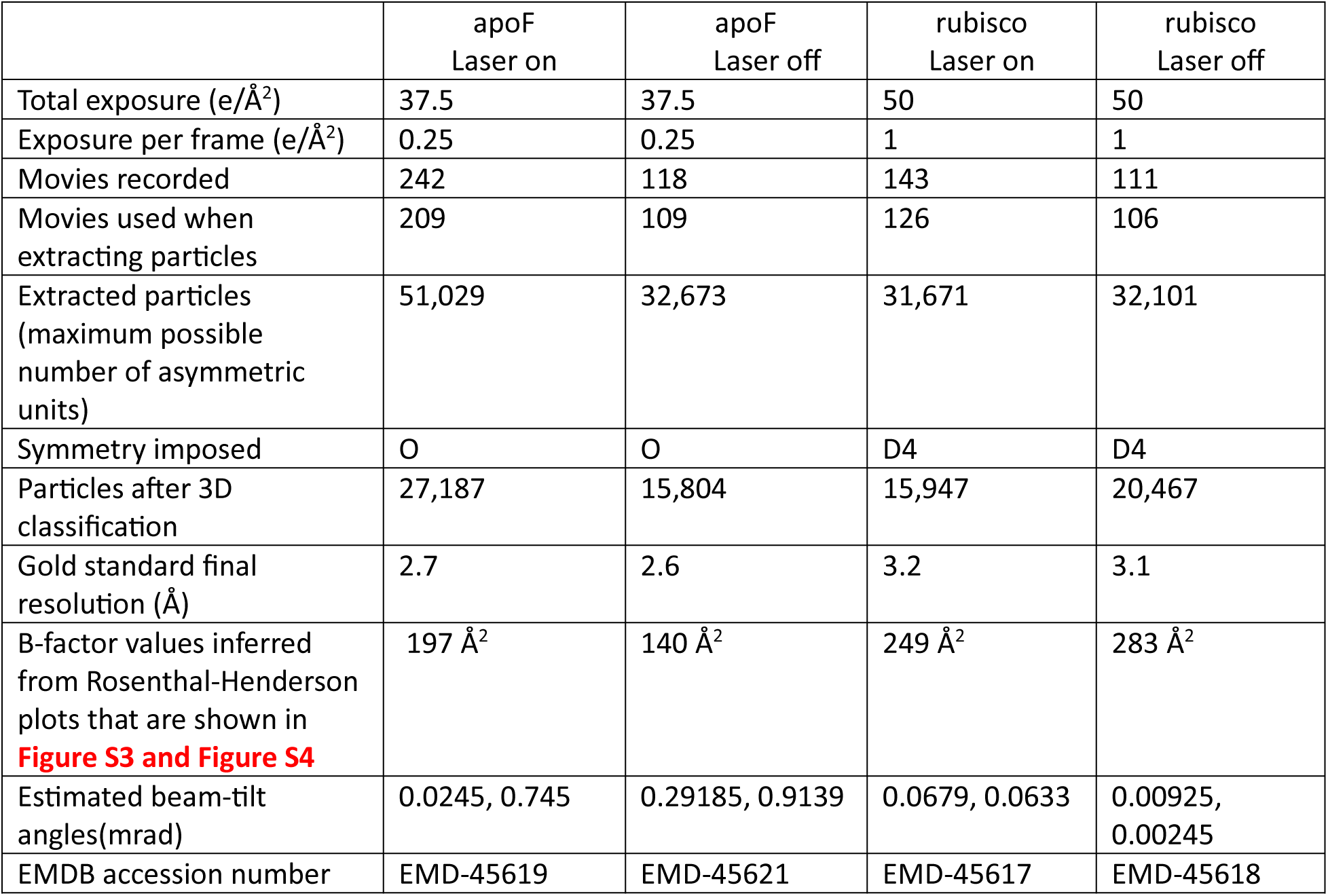
Data collection, analysis, and model-refinement statistics for images recorded in the Phase-Plate mode (relay optics turned on) of a modified Titan 80-300. Note that use of the Phase Plate mode increases the values of spherical and chromatic aberration coefficients to 4.8 mm and 7.2 mm, respectively. All images were recorded at 300 keV in movie mode, using a Gatan K2 camera and a magnification that produced a pixel size, referred to the specimen, of 0.7 Å. As indicated in the column headings for different data sets, the laser was either on, producing phase shifts shown in **Figure 1**, or off. In all cases the defocus values ranged between about 0.8 µm and 1.5 µm.

### Data analysis

Movies were imported into cryoSPARC 4.4.0 (Punjani et al., 2017), where patch motion correction was performed with dose weighting. Motion-corrected micrographs were then inspected to remove what were judged to be bad images. The most common reasons to remove images were excessive beam-induced motion and poor results of CTF fitting, observed as either low correlation with observed Thon rings at high resolution, or spurious phase-shift estimation. See Table 1 for details of the number of movies recorded and the number of movies used for processing.

CTFFIND 4.1.14 (Rohou and Grigorieff, 2015) was used to fit defocus and phase-shift values without attempting to account for the presence of the central streak in the image’s power spectra, which is due to the laser standing wave (Schwartz et al., 2019). Defocus values in these data sets typically ranged between 0.8 and 1.5 µm. **Figure 1**, which is similar to Figure 3 in (Axelrod et al., 2024), documents the degree of stability of the phase shift achieved, the value of which only rarely deviated by more than 10° from the target value of 90° during automated data collection.

**Figure 1.**
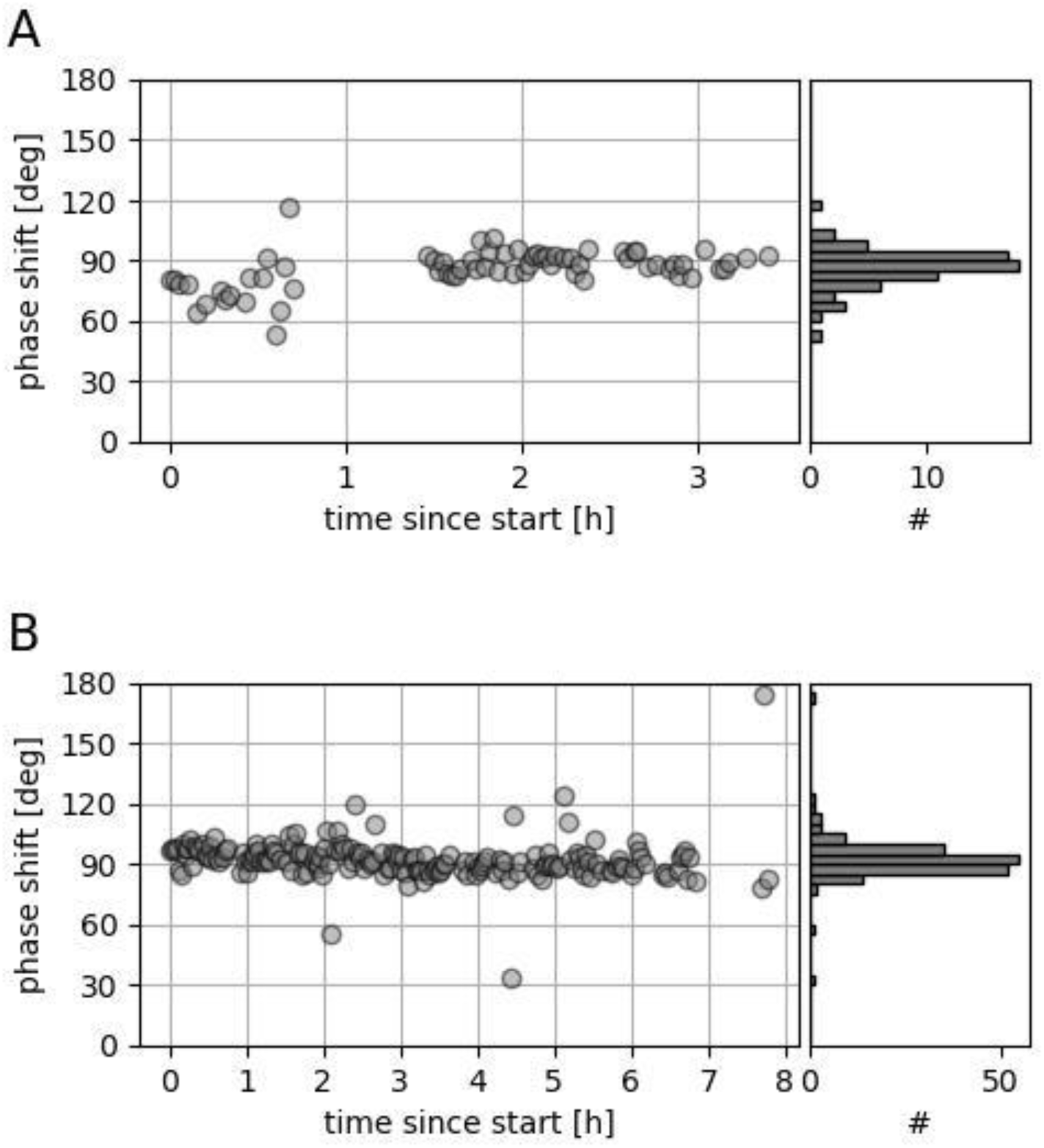
Stability of the LPP phase shift over the time-course of collecting full data sets. (A) Distribution of values for the apoF data set. A similar example of the stability of the LPP has been included as part of a recent review (Axelrod et al., 2024). (B) Corresponding values for the rubisco data set.

**Figure 2.**
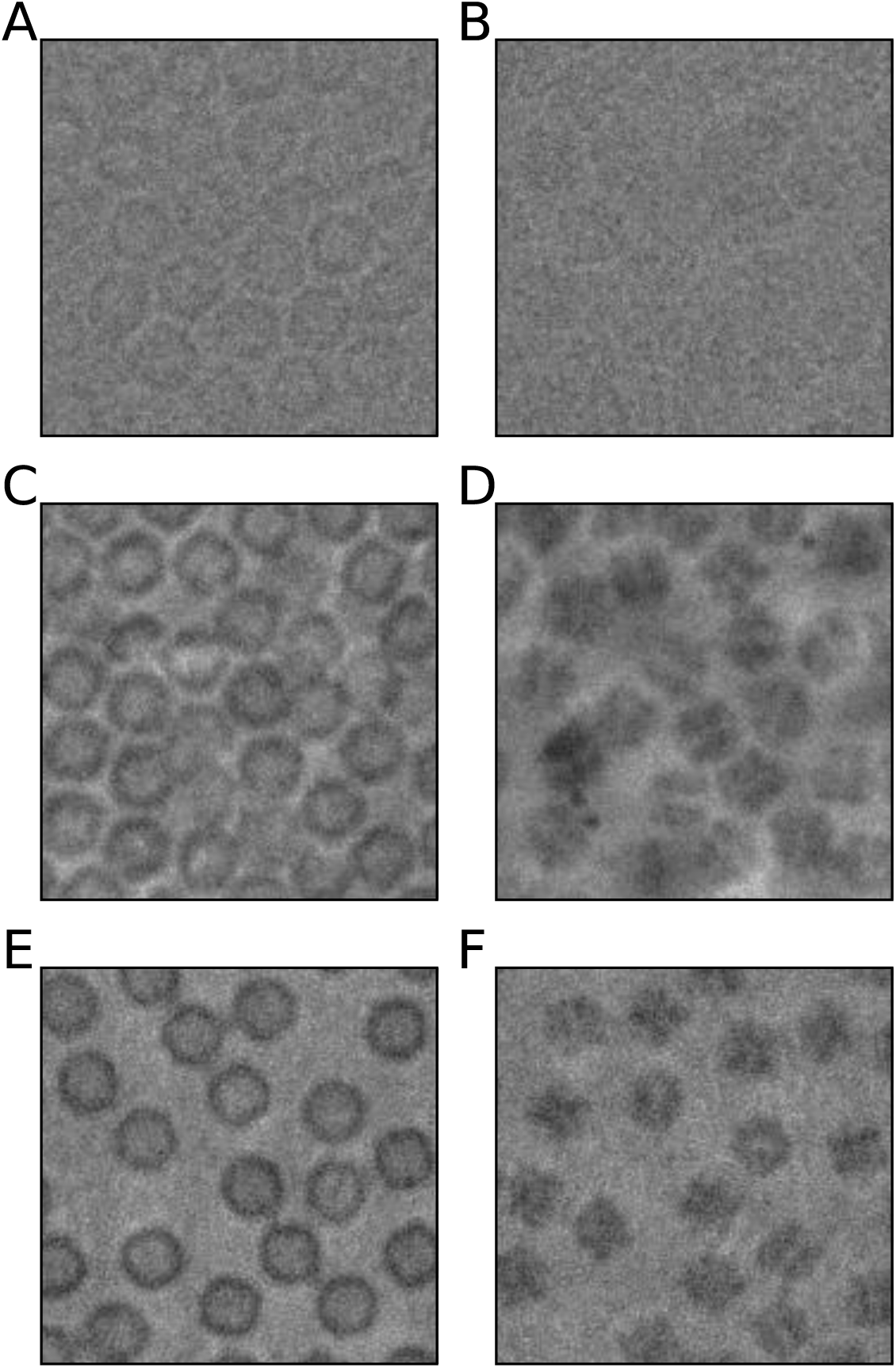
Comparison of experimental and simulated cryo-EM images. Top row: Experimental images of (A) apoF and (B) rubisco, obtained without a phase plate. Middle row: Experimental LPP images of (C) apoF and (D) rubisco. Bottom row: Simulated LPP images of (E) apoF and (F) rubisco.

**Figure 3.**
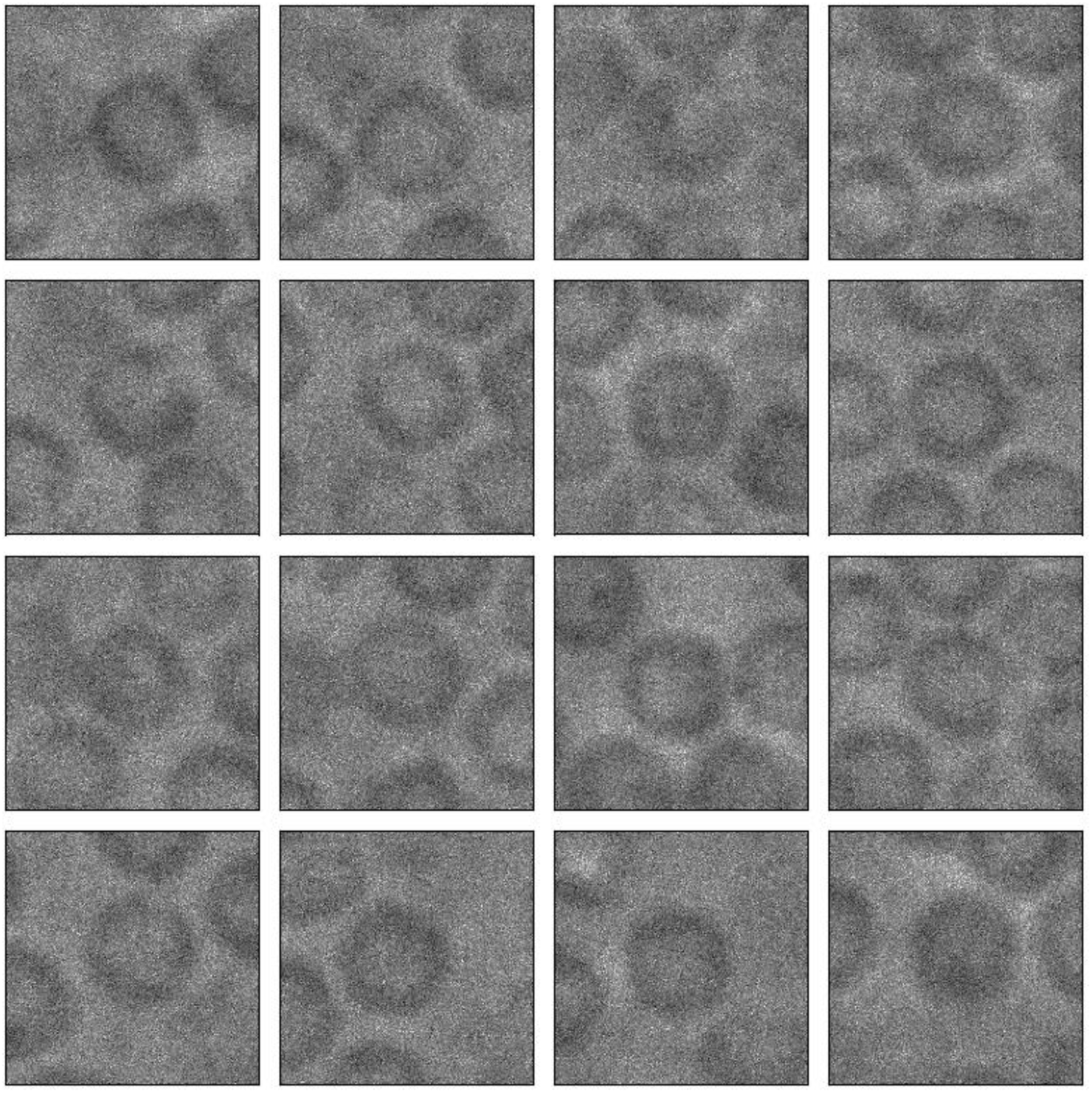
Representative examples of particle images of apoF taken from the final stack of LPP images used to obtain the refined map shown in **Figure S3**. The full stack is available at accession numbers EMPIAR-12210 (laser off) and EMPIAR-12211 (laser on), respectively.

The cryoSPARC “blob picker” program was used to pick an initial set of particles, which were extracted with a box size consisting of 384 pixels on edge. These particles were 2-D classified, and good classes were chosen as templates for reference-based particle picking. The resulting set of particles was then used for a round of 2-D classification, and good particle-classes were selected to go through a round of multi-class, *ab-Initio* reconstruction to generate an initial set of 3-D volumes. The particles that produced what was judged to be the best volume were then run through a homogeneous refinement job in cryoSPARC (Punjani et al., 2020), applying the appropriate symmetry operation and doing both Defocus and Global CTF Refinement jobs. The resultant volume and particle alignments were then used to do reference-based motion correction, implemented within cryoSPARC.

These aligned particles were exported to RELION 4.0 (Kimanius et al., 2021) using pyem (Asarnow et al., 2019). Within RELION, the particle set underwent 3D classification without image alignment, and the particles from classes displaying high-resolution features were selected and exported back into cryoSPARC. A final round of homogeneous refinement was done as well as using the map sharpening utility to generate the final reconstructed volume. These final particle sets were then used to create data for estimating the B-factor by fitting to plots of the resolution-squared *vs* number of particles (Rosenthal and Henderson, 2003), where each subset was subjected to homogeneous refinement.

### Image simulations

Python programs used for image simulation were written in-house, taking into account the modifications of the contrast transfer function (CTF) that are introduced by our current implementation of the LPP, shot noise, a model of vitreous ice, particle orientation, and radiation damage. The numerical aperture of the laser cavity was assumed to be 0.05, the electron diffraction pattern was assumed to be centered on an antinode of the laser standing wave, and no electron-optical wave aberrations other than defocus and spherical aberration were included.

Images were simulated using the atomic positions for the Protein Data Bank (PDB) models 6z6u (apoF) and 7smk (rubisco). Particles were rotated by randomly-selected Euler angles and placed at arbitrary positions within the field of view. Projected potentials were computed using atomic scattering factors parametrized as described in (Kirkland, 1998). A continuous solvation potential for each protein was computed following the procedure described by (Shang and Sigworth, 2012). The solvent was then discretized by transforming the continuous water potential into a probability density using the number density corresponding to low-density amorphous ice (∼0.0314 Å^-3^). This probability density was randomly sampled in three dimensions to generate water-molecule positions with appropriate local density. The projected potential contributed by the water molecules was approximated as that of oxygen, rescaled to produce the correct inelastic-scattering mean free path in bulk ice (Himes and Grigorieff, 2021). The (real-valued) projected potential was computed at a pixel size of 0.35 Å and converted to the corresponding phase modulation of 300-kV electrons.

The contrast transfer function (CTF) was applied as described in (Rohou and Grigorieff, 2015) to account for defocus, spherical aberration, amplitude contrast, and – in this case – the LPP phase shift described in Supplementary Note 2 of (Schwartz et al., 2019). Envelopes due to spatial coherence, temporal coherence, radiation damage (Grant and Grigorieff, 2015), and thermal magnetic field noise (Axelrod et al., 2023) were applied in the Fourier domain. In addition, the K2 camera modulation transfer function was applied, using a curve provided in a *.star file format by Gatan Inc., assuming a Nyquist frequency of 1/(1.4 Å). To take radiation damage into account, images were simulated as 50-frame stacks with an exposure of 1 e-/Å^2^/frame, and Poissonian noise was applied to each frame. Finally, images were down-sampled to a pixel size of 0.7 Å by Fourier cropping, and further binning and normalization was applied, as appropriate, to match that applied to experimental images whenever comparisons are shown. Scripts used to perform these simulations are available upon request.

## RESULTS

As is well-established, the contrast and signal-to-noise ratio at low spatial frequencies of single-particle cryo-EM images is significantly improved by using a phase plate. This point is illustrated in **Figure 2**, where examples of images recorded without a phase plate, **panels (A) and (B)**, are compared to ones recorded with the LPP, **panels (C) and (D)**. All images are shown after gain-normalization, motion-correction, dose-weighting, and down-sampling to a final pixel size of 1.4 Å, and the intensity values that are displayed span the range of ±3σ about the mean for the respective images. Full fields of view from which **panels (C) and (D)** were extracted are shown in **Figure S1**, where the extracted areas are indicated by square outlines. For comparison, full fields of view from which **panels (A) and (B)** were extracted are shown in **Figure S2**. Complete sets of the experimental images used to extract particles, together with the final particle-stacks used to produce refined density maps, are available with accession numbers EMPIAR-12208, EMPIAR-12209, EMPIAR-12210, and EMPIAR-12211.

Importantly, the increased contrast in the LPP images makes particle-to-particle heterogeneity (see examples shown in **Figure 2, C and D**) too apparent to be dismissed as being due to shot noise or differences in particle orientation. Instead, the perimeters of some apoF particles are seen to be broken or incomplete, and many other particles show non-uniform or excess density around their perimeters. In the case of rubisco, mass density (contrast) appears higher or lower in some portions of the 2D projections, depending upon the particle, than is expected for native particles in vitreous ice.

A similar degree of heterogeneity is not seen in the simulated images shown in **Figure 2E and Figure 2F**, which reflect differences that are due to shot noise and in particle orientation, but not any differences that are due to compositional variation or damage. We believe that the differences between these experimental and simulated LPP images are too great to be reasonably explained by native conformational differences, which usually involve only small movements of flexible subunits. We therefore infer that many particles are damaged to an extent that had not previously been appreciated.

We thus asked whether experimental images of the type shown in **Figure 2** would produce high-resolution maps, taking into consideration the limitations of our modified Titan electron microscope, (Axelrod et al., 2023). Parenthetically, coma and 3-fold astigmatism also emerged as potentially resolution-limiting effects during processing of the apoF data sets used here. This discovery required that further steps be taken when aligning the electron beam to the laser beam in the LPP, which were added to our automated protocol when collecting the rubisco data set. These aberrations have no effect on the SNR at low resolution, of course, which is the issue addressed here.

Despite the limitations of our modified Titan, the resolution obtained from 27,187 apoF particles was 2.7 Å when using the LPP, while the resolution improved to 2.6 Å, from only 15,804 apoF particles, when the laser was turned off (but with the electron relay optics still turned on), as is summarized in **Table 1**. Possible reasons for the diminished performance – when the laser was turned on – are suggested in the Discussion section. At the same time, when imaged in “standard mode” of the same microscope, i.e. with the relay optics turned off, the resolution obtained from about 6000 apoF particles improved to 2.0 Å, as previously reported in Figure 2 of (Axelrod et al., 2023). This latter result confirms our expectation that the electron relay optics, as currently implemented, limit the resolution to a greater extent than does the particle quality. In the case of rubisco, 15,947 laser-on particle-images result in a 3.2 Å reconstruction while 20,467 laser-off particle-images results in a 3.1 Å reconstruction. Additional documentation supporting these results is presented as **Figure S3 and Figure S4.**

Perhaps surprisingly, considerable amounts of apparent heterogeneity persist in many of the particle images that are included in the final stacks used to obtain high-resolution reconstructions. **Figure 3** shows examples of particles taken randomly from the final stack of apoF images, while **Figure 4** shows corresponding examples taken from the final stack of rubisco particles. The unexpected observation that images of many “damaged” particles appear to remain in the final stacks raises the question of how much value might be gained by first removing them, perhaps by using the approached described in (Zhu et al., 2023). We consider this question to be separate from the main issue presented here, however, which is that the LPP has made it much easier to detect such heterogeneity. At the same time, this and other issues can also be investigated by using the full data sets that have been deposited.

**Figure 4.**
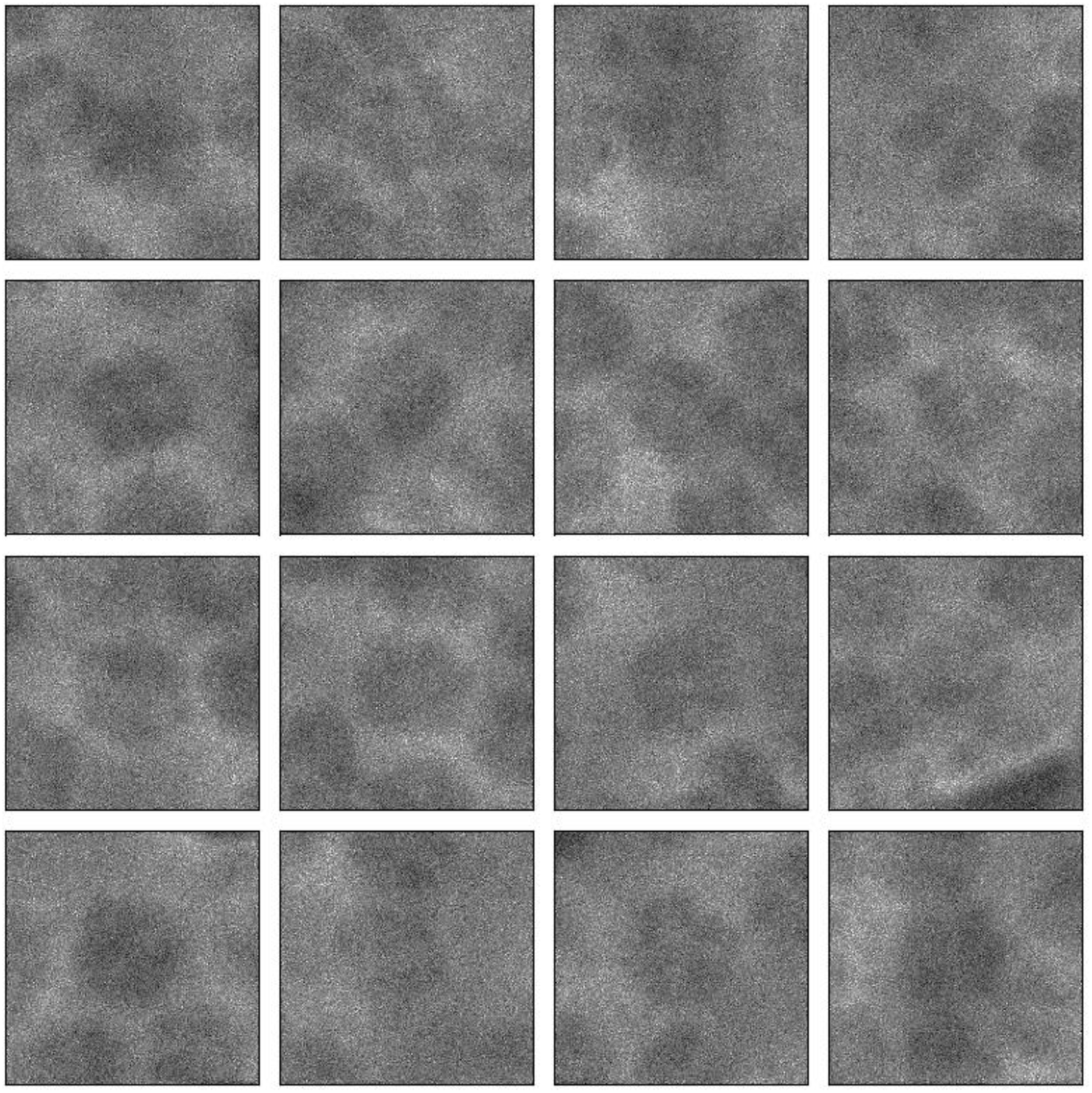
Representative examples of particle images of rubisco taken from the final stack of LPP images used to make the refined map shown in **Figure S4**. The full stack is available at accession numbers EMPIAR-12208 (laser off) and EMPIAR-12209 (laser on), respectively.

## DISCUSSION

### Image contrast is greatly increased at low spatial frequencies

The basis for increased contrast at low frequencies is illustrated in **Figure 5**, where the modulus of the contrast transfer function (CTF) of our current LPP is compared to that for defocus alone. The legend for **Figure 5** provides further information about the parameters used to calculate the respective CTF curves, including the fact that a defocus value of 1 µm was assumed in both cases. We note that Thon rings are currently used to accurately estimate the actual defocus values in low-dose images, which frequently differ from the intended defocus values, and thus a defocus of few hundred nm or more must be used even with phase plates (Danev et al., 2017).

**Figure 5.**
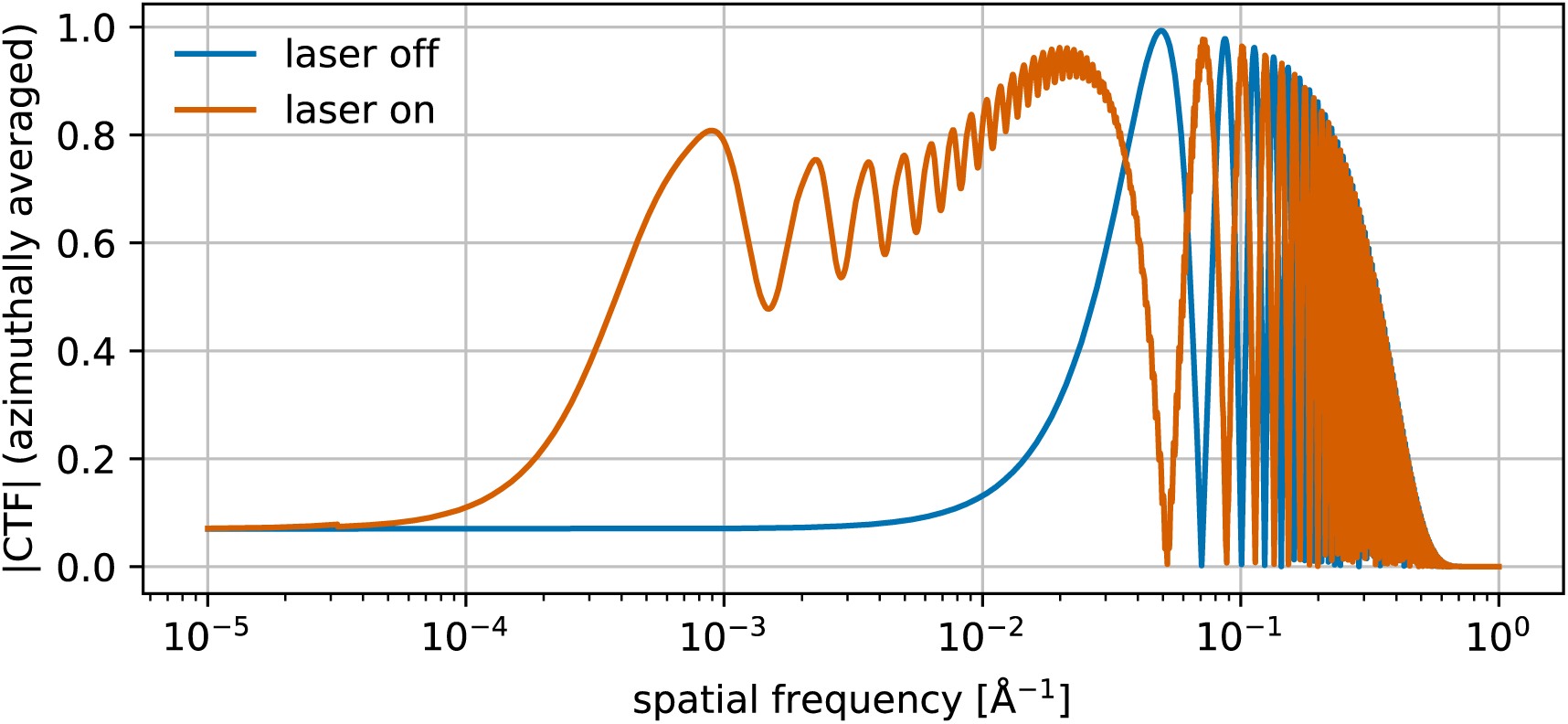
Radially averaged modulus of CTF curves for our modified Titan electron microscope, when operated with its relay optics turned on. The CTF when a 90° phase shift is applied to the unscattered electron beam is shown by the orange curve, while the CTF due to defocus only, i.e. when the laser is off, is shown in blue. The following instrumental parameters apply to both curves: amplitude contrast 0.07, underfocus 1 µm, Cs 4.8 mm, Cc 7.6 mm, untilted (parallel) illumination, and 0.137 Å^2^ Johnson noise (Axelrod et al., 2023). The LPP phase shift produced by the laser standing wave is described in Supplementary Note 2 of (Schwartz et al., 2019). Neither curve includes the effect of the envelope function due to a camera MTF.

Since the spectral power of images is proportional to the square of the CTF, the signal power of LPP images is expected to exceed that for images obtained with defocus alone by a factor of 10 to 100 over the range of spatial frequencies from ∼1/(2500 Å) to ∼1/(50 Å). As a result, it is understandable that visual detection of per-particle structural heterogeneity, examples of which were shown in **Figure 2**, **Figure 3, and Figure 4**, is greatly improved when using the LPP. Conversely, this heterogeneity has gone largely unnoticed for conventional cryo-EM images, which are usually recorded with as little defocus as feasible.

It remains to be seen whether the signal at frequencies higher than ∼1/(50 Å) can also be recovered more completely when using a phase plate. Doing so will require developing a method, other than using Thon rings, to experimentally acquire and computationally process images that are “close-to-focus”, thereby avoiding the oscillations in the CTF that set in at frequencies as low as ∼1/(20 Å). Using such in-focus images without a phase plate is not a realistic option, of course, since such images would only exhibit amplitude contrast, the magnitude of which is but a small fraction of that for phase contrast.

### The increased signal of LPP images is best appreciated after standard contrast adjustment

The per-particle heterogeneity, reported here, is still difficult to recognize by eye in unprocessed LPP images, such as those shown in **Figure S5**. Instead, per-particle heterogeneity becomes more readily apparent after binning and normalizing, which are standard operations used during single-particle data processing (Scheres et al., 2009; Sorzano et al., 2004). Heterogeneity becomes visually quite clear, for example, when the displayed image intensity covers the reduced range of ±3σ about the mean value, as is shown in **Figure 2**, **Figure 3, and Figure 4**.

Even after contrast adjustment, however, the same heterogeneity was not as readily noticed in conventional cryo-EM images of the same specimens, which were recorded without the use of the LPP and at the commonly-used defocus values between 0.5 µm and 1.5 µm. Nevertheless, when examining such images more critically, similar heterogeneity is also seen in our “laser off” data sets after binning and normalization, (see **Figure S2**), albeit at lower signal-to-noise ratio (SNR) than that which is produced with the LPP. Because of the increase signal that is provided in LPP images at low resolution, however, this heterogeneity can no longer be ignored, as is shown in **Panels C and D of Figure 2** (and annotated further in **the graphical abstract**).

Significantly, the same degree of heterogeneity is not seen in the case of simulated LPP images shown in **Panels E and F of Figure 2**. In this latter, ideal case, only much smaller particle-to-particle variations are modeled, i.e. those due to orientation-dependent differences in the projected structure and those due to shot noise.

### Some trivial-artifacts can be ruled out as causing heterogeneity

Simulations rule out the possibility that the heterogeneity seen in **Panels C and D of Figure 2** might be a feature that is produced by the LPP. Simulated images do show two image artifacts that are expected for LPP images – “halos”, also present in images produced with the VPP (Danev et al., 2014), and “ghost images”, uniquely generated by the LPP (Schwartz et al., 2019). Both of these features are quite faint, however, as is demonstrated by the simulated images shown in **Panels E and F of Figure 2**. While effects due to inelastically scattered electrons, specimen motion, and misalignment of the laser beam to the electron beam were not included as part of the simulations, none of these would provide reasonable explanations for the apparent heterogeneity seen in the experimental images. As a result, we believe that the main cause of discrepancy between the simulated and experimental images shown in **Figure 2** must be attributed to features in the vitrified specimens, which are not present in our model of vitreous ice or in the PDB models used in the simulations.

We further rule out the suggestion that the particle heterogeneity seen in our experimental LPP images is unique to our specimen grids. On the contrary, similar levels of structural heterogeneity can also be seen in enlarged versions of apoferritin images that were recorded with a VPP, for example Figure 3B of (Li et al., 2019) or Figure 1C of (Fan et al., 2017). Indeed, similar amounts of heterogeneity can also be seen – once one begins to look for it – in Extended Data Figure 3b of (Küçükoğlu et al., 2024). This latter figure shows an example of the CTEM images that were used to obtain an apoF map at a resolution of 1.09 Å. Furthermore, small spots with high mass density (perhaps e.g. protein fragments) and faint-contrast particles are apparent in VPP images of the 20S proteasome, which are shown as Figure 1A and Figure 2A of (Danev et al., 2017).

### Heterogeneity thus appears to reflect structural damage

It is well established that adsorption of biological macromolecules to the air-water interface (AWI) often results in preferential particle-orientation (Carragher et al., 2019; Drulyte et al., 2018; Noble et al., 2018) and – importantly – even severe structural damage (D’Imprima et al., 2019). It remains uncertain, however, whether the damage revealed here, by using the LPP, first occurs following contact with the air-water interface (AWI), or whether it was already there to begin with, while the particles were still in the test tube.

The insight gained by using a phase plate leads us to speculate that similar heterogeneity may affect a large fraction of the particles that are extracted from images of most types of particles. Evidence supporting this view includes the fact that many candidate particles are rejected during 2-D and 3-D classification, when extracted particles fall into classes that traditionally are regarded as noise or “junk”. In addition, it has recently been discovered that even a large fraction of the particle images in the final stacks that are used to produce high-resolution density maps actually contribute little, if anything, to the final map (Zhu et al., 2023).

### High resolution is achieved in spite of many extracted particles being visibly damaged

Despite the particle-heterogeneity that is easily revealed when using the LPP, high-resolution structures are obtained from such specimens when symmetry is enforced, as is documented in **Figure S3 and Figure S4**. Indeed, we obtained maps consistent with that of a high-quality sample, judged by conventional standards, when using our grids to collect images of apoF, both in our low-base Titan microscope without using the relay optics (Figure 2 of (Axelrod et al., 2023), as well as with another microscope for rubisco (SI Figure 4 of (Blikstad et al., 2023)). Given the limitations that can be expected from our low-base Titan, which is equipped with only a side-entry cold stage, the results presented here compare not too unfavorably to the higher-resolution results that others have obtained with better microscopes (Küçükoğlu et al., 2024; Nakane et al., 2020; Yip et al., 2020; Zhang et al., 2020).

The resolution that is achieved in the current work takes on further significance in light of the fact that even the subsets of particles used to produce symmetrized maps are quite heterogeneous, as is illustrated in **Figure 3 and Figure 4**. It thus seems apparent that previously unrecognized structural heterogeneity, the amount of which varies randomly from one particle to the next, leaves sufficiently large portions of the initially symmetrical protein complexes in a state that can still be merged to produce symmetrized, high-resolution maps.

As has been described previously (Axelrod et al., 2024), increased spherical aberration, Cs, and chromatic aberration, Cc, along with a remaining contribution of Johnson noise, are expected to be the main causes for the reduced resolution that is currently achieved with the LPP. These undesirable features are expected to be reduced significantly in next-generation versions of LPP microscopes.

### Additional factors that are worth noting

Images recorded with the LPP also reveal the existence of an unexpectedly mottled background (see the **graphical abstract**, and also **Figure 2**). Although this low-frequency variation in background intensity is not apparent in images recorded without a phase plate, it can again be seen in published images that were recorded with the VPP. A denatured-protein “skin” that is formed at the AWI by apoferritin, rubisco, and many other types of proteins – see (Han and Glaeser, 2021) – might be one possible origin for the patchy variations in background. An alternative possibility is that there are significant, local variations in the thickness values of the vitrified sample.

The increased SNR at low resolution that is provided by phase plates has some potential disadvantages, such as increasing the influence that structural noise has during single-particle analysis. Structural noise is unlike shot noise in the sense that it is made more visible when using a phase plate, exactly as is the signal from undamaged portions of the particles. Among other things, the increased SNR at low resolution may thus make it especially important to avoid normalization errors when using maximum likelihood software for SPA – for more on this issue, see the second paragraph of the Discussion section of (Scheres et al., 2009). It remains to be investigated, however, whether the resulting increase in structural noise is responsible for the diminished performance that we report here for apoF, but somewhat less so for rubisco. Also remaining to be investigated is whether accounting for the laser streak as part of the CTF correction, during refinement, might further improve the LPP performance.

## CONCLUSIONS

Use of a phase plate in cryo-EM produces a significant increase in the signal at low resolution. The corresponding increase in SNR makes it possible to recognize levels of particle heterogeneity that previously had gone undetected, as well as increasing unwanted structural noise and revealing unexpected variations in the background. Mitigating these sources of structural noise may require improved methods of sample preparation as well as introducing software that better compensates for structural noise.

The increase in (low-resolution) signal provided by the LPP is expected to have obvious benefits, of course, such as making small particles much easier to recognize, or making it possible to directly visualize flexible subunits in individual images of larger particles. The potential ability to visualize individual “floppy” subunits, which effectively disappear (i.e. they are “averaged out”) when data are merged from multiple images, may thus produce biologically informative information.

Still to be determined is whether the enhanced SNR at low spatial frequencies can improve the mitigation of beam-induced motion (BIM), which currently remains a limiting factor for highly tilted specimens. Use of the LPP, by enabling sample structures themselves to serve as “fiducials”, may reduce the electron exposures need to align successive frames by a factor of 5 or more, which might result in significant reduction in the amount of BIM that occurs within frames. Such “super-fractionated” electron exposures could have important applications in cryo-tomography as well as in single-particle cryo-EM of specimens that are tilted to high angles because the particles had adopted highly preferred orientations.

## ACKNOWLEDGEMENTS

This project was supported by the U.S. National Institutes of Health (Grant No. 5 R01 GM126011), Chan Zuckerberg Initiative (award number 2021-234606), Gordon and Betty Moore Foundation (Grant No. 9366), and a cooperative research and development agreement (CRADA) with ThermoFisher Scientific (award number AWD00004352). Grant No. 5 R01 GM126011 and award number AWD00004352 were administered at Lawrence Berkeley National Laboratory under Contract No. DE-AC02-05CH11231. We thank Prof. David Savage and Dr. Luke Oltragge for providing the samples of rubisco that were used in this work. P.N.P. acknowledges support from a postdoctoral fellowship from the National Institute of General Medical Sciences of the National Institutes of Health under Award Number F32GM149186.

## SUPPLEMENTAL MATERIAL

**Figure S1.**
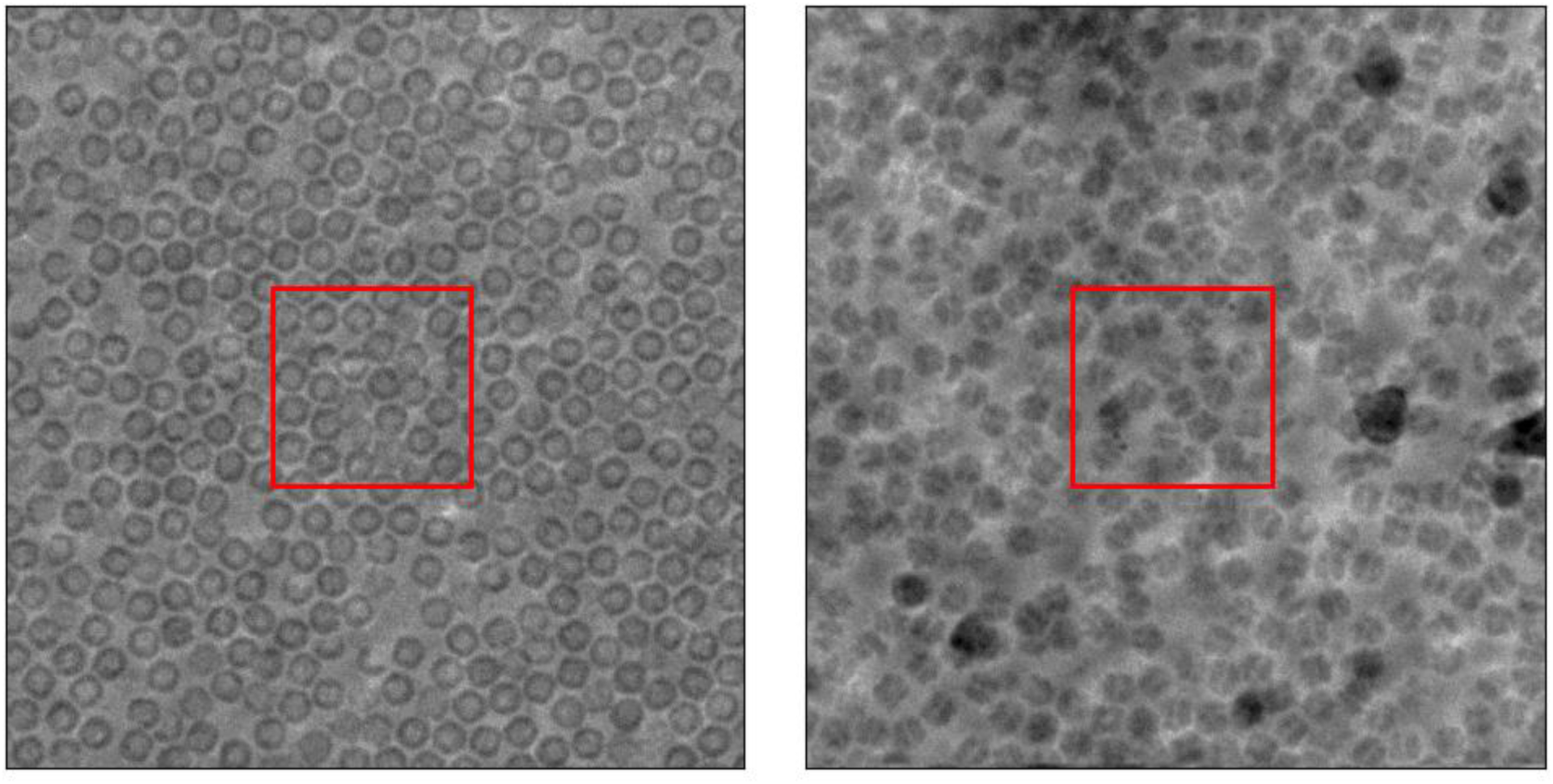
Examples of images of apoferritin (left) and rubisco (right) that were obtained when using the laser phase plate (LPP). The square areas outlined in red are shown in the main text as **Figure 2C and Figure 2D**, respectively.

**Figure S2.**
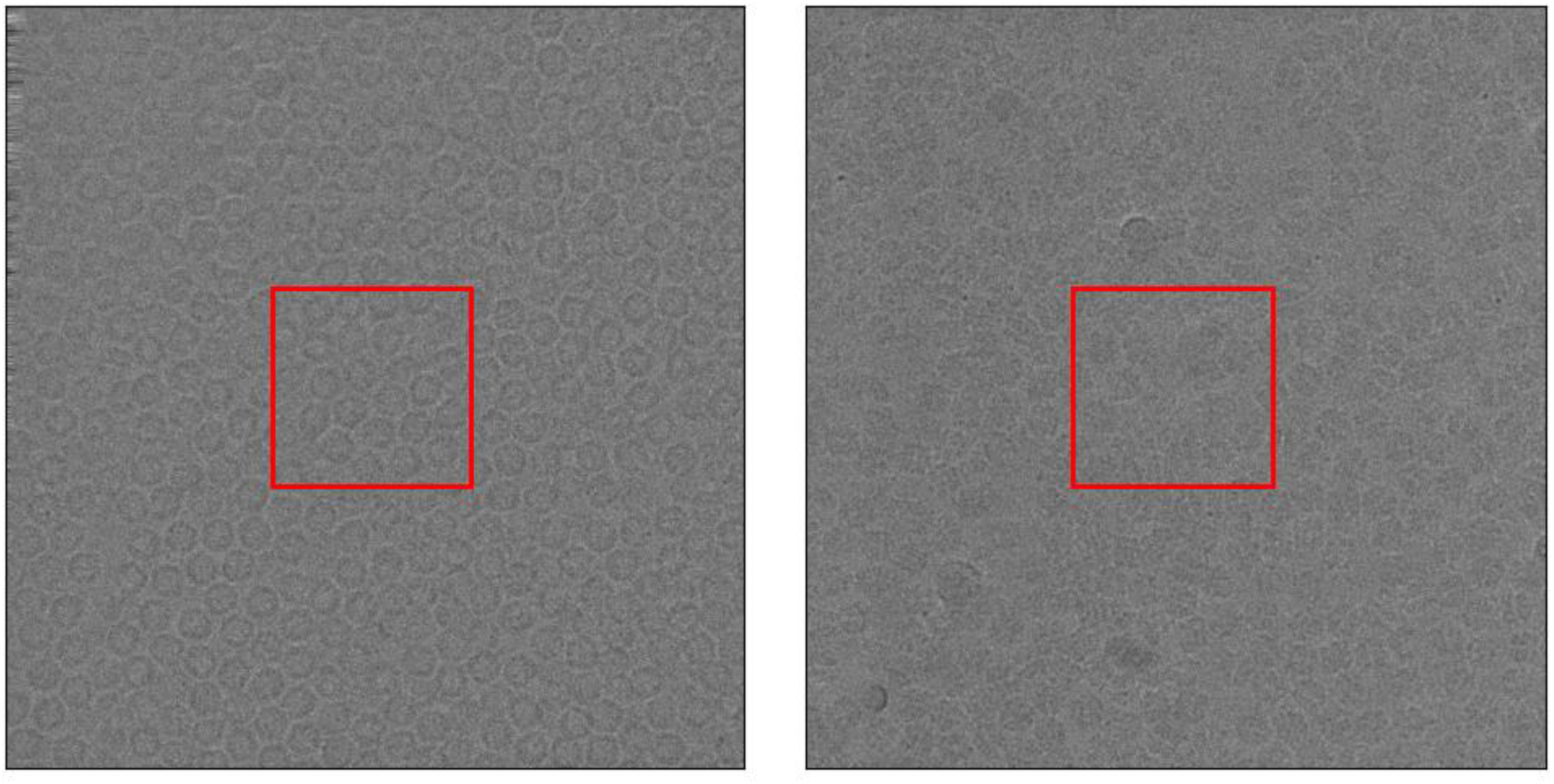
Examples of images of apoferritin (left) and rubisco (right) that were obtained when the laser was not turned on. The square areas outlined in red are shown in the main text as **Figure 2A and Figure 2B**, respectively.

**Figure S3.**
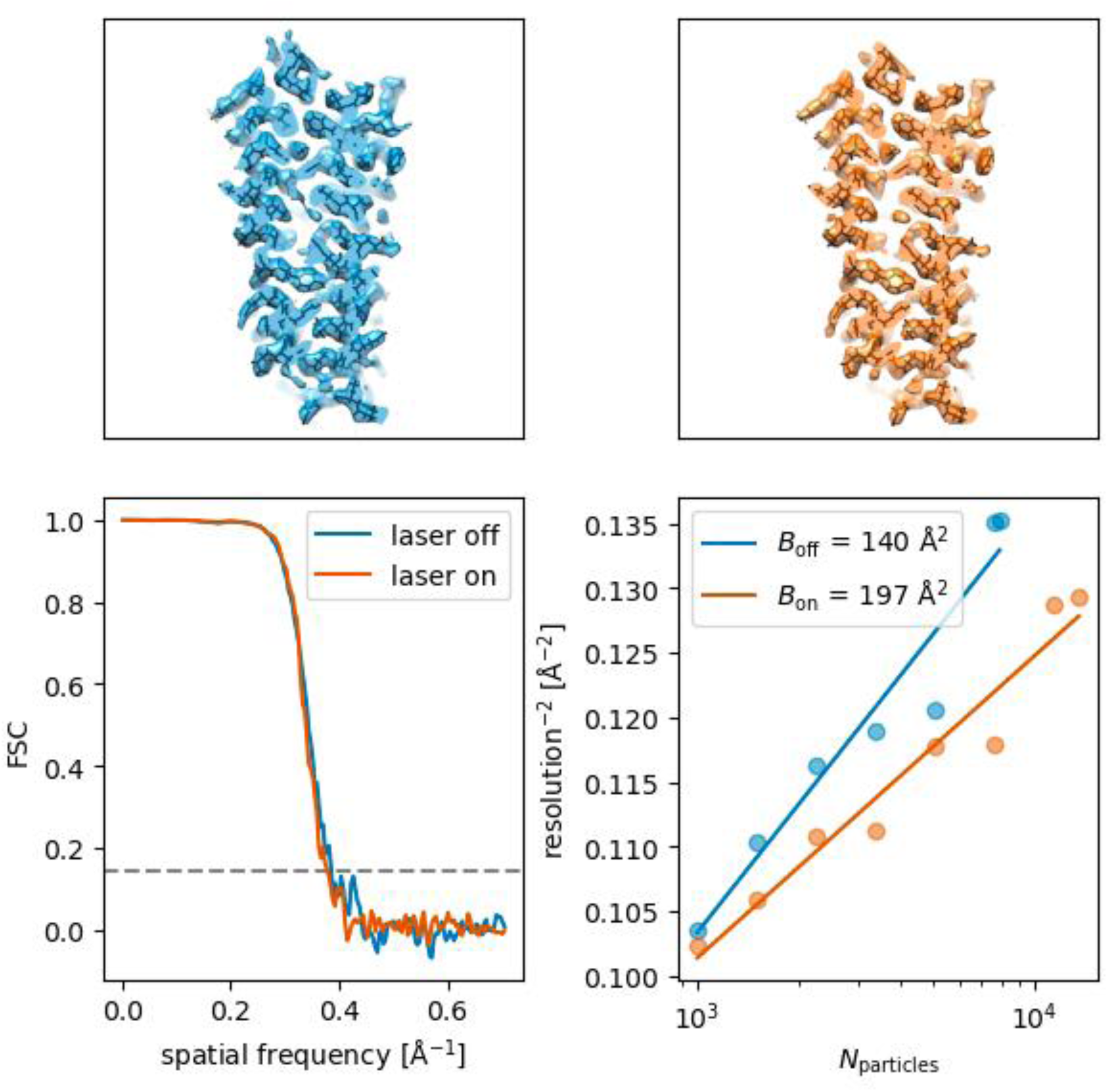
Comparison of metrics for the apoferritin maps generated from images obtained with the relay optics of our modified Titan turned on, and with the laser either turned on (indicated in orange) or turned off (indicated in blue). As is shown here, and as is noted in **Table 1**, the B-factor for apoferritin was larger for the data set obtained with the laser on, and thus a larger number particles had to be used to reach a similar resolution.

**Figure S4.**
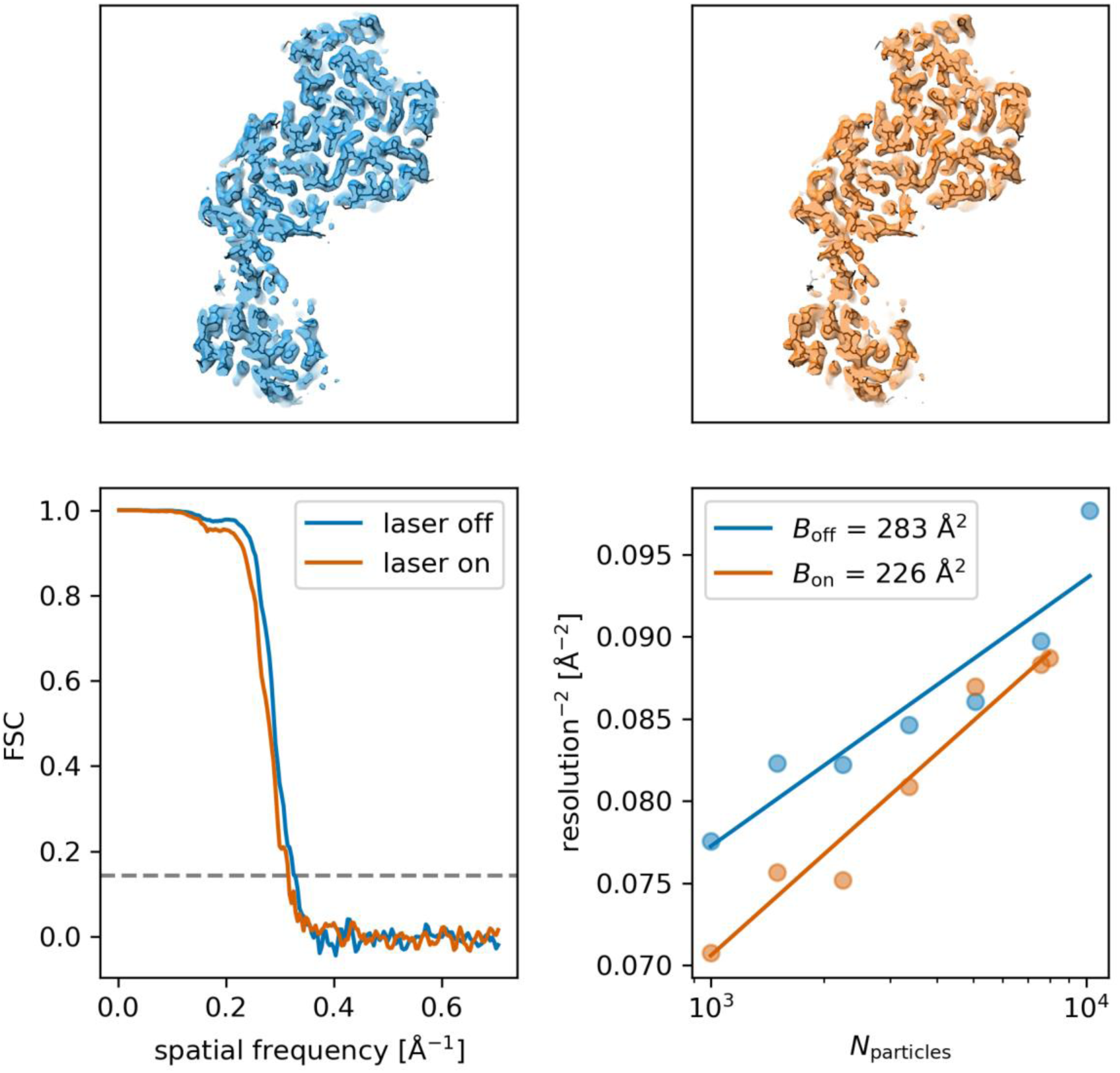
Comparison of metrics for the rubisco maps generated from images obtained with the relay optics of our modified Titan turned on, and with the laser either turned on (indicated in orange) or turned off (indicated in blue). Although the straight-line fits to the Rosenthal-Henderson plots do not extrapolate to the same values at low numbers of particles, the slopes (B-factors) are similar.

**Figure S5.**
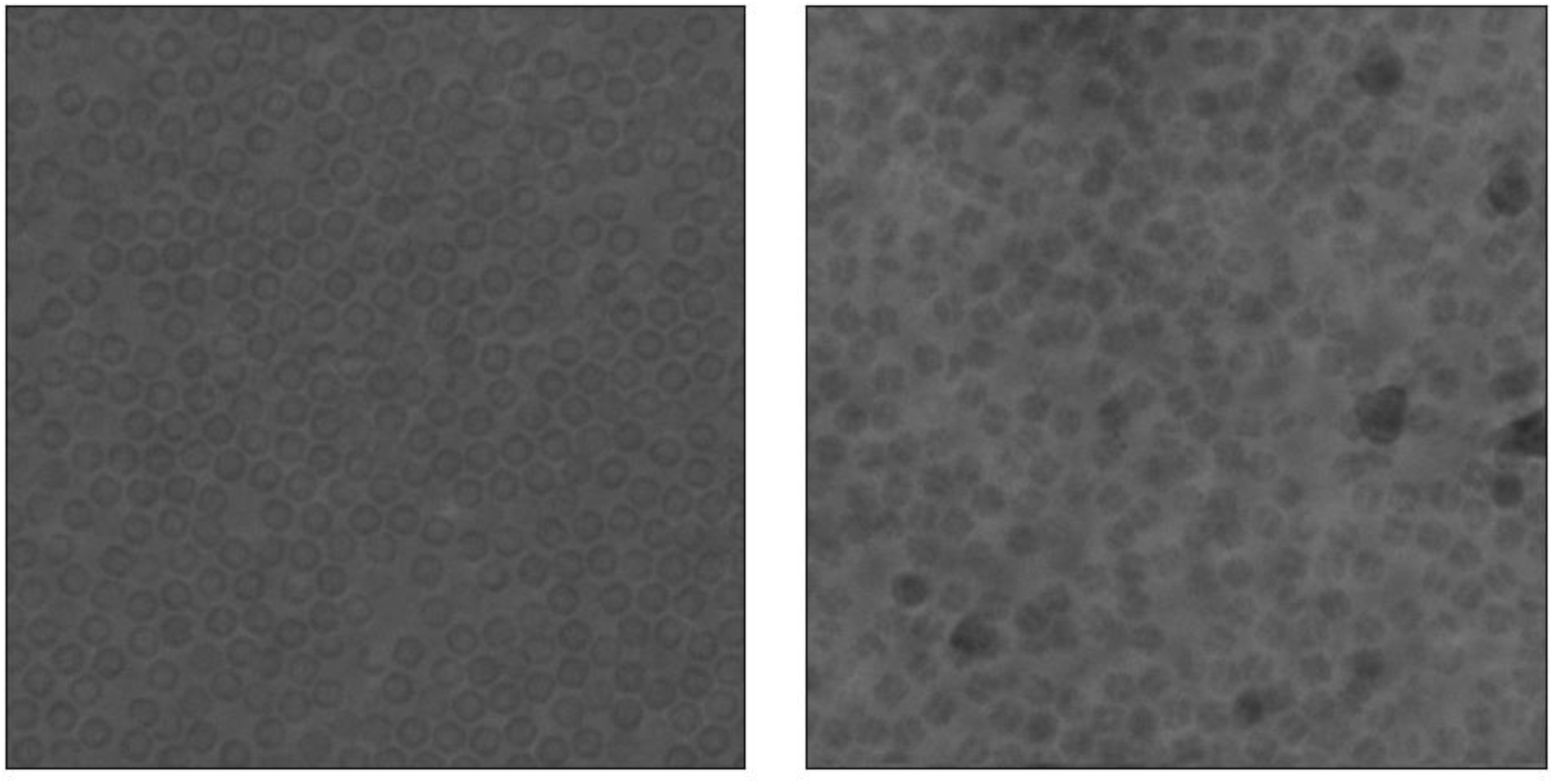
The same images of apoferritin (left) and rubisco (right) that are shown in **Figure S1**, displayed without contrast enhancement provided by binning and normalization (and without the red square).

## REFERENCES

1. Asarnow, D., E. Palovcak, Y. Cheng, 2019. pyem v0.5. Zenodo.

2. Axelrod, J.J., J.T. Zhang, P.N. Petrov, R.M. Glaeser, H. Müller, 2024. Modern approaches to improving phase contrast electron microscopy. Current Opinion in Structural Biology 86, 102805.

3. Axelrod, J.J., P.N. Petrov, J.T. Zhang, J. Remis, B. Buijsse, R.M. Glaeser, H. Mȕller, 2023. Overcoming resolution loss due to thermal magnetic field fluctuations from phase plates in transmission electron microscopy. Ultramicroscopy 249, 113730.

4. Baxter, W.T., R.A. Grassucci, H. Gao, J. Frank, 2009. Determination of signal-to-noise ratios and spectral SNRs in cryo-EM low-dose imaging of molecules. Journal of Structural Biology 166, 126–132.

5. Blikstad, C., E.J. Dugan, T.G. Laughlin, J.B. Turnšek, M.D. Liu, S.R. Shoemaker, N. Vogiatzi, J.P. Remis, D.F. Savage, 2023. Identification of a carbonic anhydrase–Rubisco complex within the alpha-carboxysome. Proceedings of the National Academy of Sciences 120, e2308600120.

6. Campbell, M.G., A.C. Cheng, A.F. Brilot, A. Moeller, D. Lyumkis, D. Veesler, J.H. Pan, S.C. Harrison, C.S. Potter, B. Carragher, N. Grigorieff, 2012. Movies of Ice-Embedded Particles Enhance Resolution in Electron Cryo-Microscopy. Structure 20, 1823–1828.

7. Carragher, B., Y. Cheng, A. Frost, R.M. Glaeser, G.C. Lander, E. Nogales, H.W. Wang, 2019. Current outcomes when optimizing ‘standard’ sample preparation for single-particle cryo-EM. Journal of Microscopy 276, 39–45.

8. D’Imprima, E., D. Floris, M. Joppe, R. Sánchez, M. Grininger, W. Kühlbrandt, 2019. Protein denaturation at the air-water interface and how to prevent it. Elife 8, e42747.

9. Danev, R., D. Tegunov, W. Baumeister, 2017. Using the Volta phase plate with defocus for cryo-EM single particle analysis. Elife 6, e23006.

10. Danev, R., B. Buijsse, M. Khoshouei, J.M. Plitzko, W. Baumeister, 2014. Volta potential phase plate for in-focus phase contrast transmission electron microscopy. Proceedings of the National Academy of Sciences 111, 15635–15640.

11. Drulyte, I., R.M. Johnson, E.L. Hesketh, D.L. Hurdiss, C.A. Scarff, S.A. Porav, N.A. Ranson, S.P. Muench, R.F. Thompson, 2018. Approaches to altering particle distributions in cryo-electron microscopy sample preparation. Acta Crystallographica Section D-Structural Biology 74, 560–571.

12. Fan, X., L. Zhao, C. Liu, J.-C. Zhang, K. Fan, X. Yan, H.-L. Peng, J. Lei, H.-W. Wang, 2017. Near-Atomic Resolution Structure Determination in Over-Focus with Volta Phase Plate by Cs-Corrected Cryo-EM. Structure 25, 1623–1630.e3.

13. Glaeser, R.M., 2013. Invited Review Article: Methods for imaging weak-phase objects in electron microscopy. Review of Scientific Instruments 84, 111101.

14. Glaeser, R.M., E. Nogales, W. Chiu, 2021. Single-particle Cryo-EM of Biological Macromolecules, in: Glaeser, R M, et al., Eds.), IOP Publishing.

15. Grant, T., N. Grigorieff, 2015. Measuring the optimal exposure for single particle cryo-EM using a 2.6 Å reconstruction of rotavirus VP6. Elife 4, e06980.

16. Han, B.-G., R.M. Glaeser, 2021. Simple assay for adsorption of proteins to the air–water interface. Journal of Structural Biology 213, 107798.

17. Himes, B., N. Grigorieff, 2021. Cryo-TEM simulations of amorphous radiation-sensitive samples using multislice wave propagation. IUCrJ 8, 943–953.

18. Kimanius, D., L. Dong, G. Sharov, T. Nakane, S.H.W. Scheres, 2021. New tools for automated cryo-EM single-particle analysis in RELION-4.0. Biochem J 478, 4169–4185.

19. Kirkland, E.J., 1998. Advanced computing in electron microscopy. Plenum Press.

20. Küçükoğlu, B., I. Mohammed, R.C. Guerrero-Ferreira, S.M. Ribet, G. Varnavides, M.L. Leidl, K. Lau, S. Nazarov, A. Myasnikov, C. Sachse, K. Müller-Caspary, C. Ophus, H. Stahlberg, 2024. Low-dose cryo-electron ptychography of proteins at sub-nanometer resolution. bioRxiv, 2024.02.12.579607.

21. Li, K., C. Sun, T. Klose, J. Irimia-Dominguez, F.S. Vago, R. Vidal, W. Jiang, 2019. Sub-3 Å apoferritin structure determined with full range of phase shifts using a single position of volta phase plate. Journal of Structural Biology 206, 225–232.

22. Mastronarde, D.N., 2005. Automated electron microscope tomography using robust prediction of specimen movements. Journal of Structural Biology 152, 36–51.

23. Nakane, T., A. Kotecha, A. Sente, G. McMullan, S. Masiulis, P.M.G.E. Brown, I.T. Grigoras, L. Malinauskaite, T. Malinauskas, J. Miehling, T. Uchański, L. Yu, D. Karia, E.V. Pechnikova, E. de Jong, J. Keizer, M. Bischoff, J. McCormack, P. Tiemeijer, S.W. Hardwick, D.Y. Chirgadze, G. Murshudov, A.R. Aricescu, S.H.W. Scheres, 2020. Single-particle cryo-EM at atomic resolution. Nature 587, 152–156.

24. Noble, A.J., V.P. Dandey, H. Wei, J. Braschi, J. Chase, P. Acharya, Y.Z. Tan, Z.N. Zhang, L.Y. Kim, G. Scapin, M. Rapp, E.T. Eng, W.J. Rice, A.C. Cheng, C.J. Negro, L. Shapiro, P.D. Kwong, D. Jeruzalmi, A. des Georges, C.S. Potter, B. Carragher, 2018. Routine single particle cryoEM sample and grid characterization by tomography. Elife 7, e34257.

25. Oltrogge, L.M., T. Chaijarasphong, A.W. Chen, E.R. Bolin, S. Marqusee, D.F. Savage, 2020. Multivalent interactions between CsoS2 and Rubisco mediate α-carboxysome formation. Nat Struct Mol Biol 27, 281–287.

26. Punjani, A., H. Zhang, D.J. Fleet, 2020. Non-uniform refinement: adaptive regularization improves single-particle cryo-EM reconstruction. Nature Methods 17, 1214–1221.

27. Punjani, A., J.L. Rubinstein, D.J. Fleet, M.A. Brubaker, 2017. cryoSPARC: algorithms for rapid unsupervised cryo-EM structure determination. Nature Methods 14, 290.

28. Rohou, A., N. Grigorieff, 2015. CTFFIND4: Fast and accurate defocus estimation from electron micrographs. Journal of Structural Biology 192, 216–221.

29. Rosenthal, P.B., R. Henderson, 2003. Optimal determination of particle orientation, absolute hand, and contrast loss in single-particle electron cryomicroscopy. Journal of Molecular Biology 333, 721–745.

30. Scheres, S.H.W., M. Valle, P. Grob, E. Nogales, J.-M. Carazo, 2009. Maximum likelihood refinement of electron microscopy data with normalization errors. Journal of Structural Biology 166, 234–240.

31. Schwartz, O., J.J. Axelrod, S.L. Campbell, C. Turnbaugh, R.M. Glaeser, H. Muller, 2019. Laser phase plate for transmission electron microscopy. Nature Methods 16, 1016–2020.

32. Shang, Z., F.J. Sigworth, 2012. Hydration-layer models for cryo-EM image simulation. Journal of Structural Biology 180, 10–16.

33. Sorzano, C.O.S., L.G. de la Fraga, R. Clackdoyle, J.M. Carazo, 2004. Normalizing projection images: a study of image normalizing procedures for single particle three-dimensional electron microscopy. Ultramicroscopy 101, 129–138.

34. Turnbaugh, C., J.J. Axelrod, S.L. Campbell, J.Y. Dioquino, P.N. Petrov, J. Remis, O. Schwartz, Z. Yu, Y. Cheng, R.M. Glaeser, H. Mueller, 2021. High-power near-concentric Fabry–Perot cavity for phase contrast electron microscopy. Review of Scientific Instruments 92, 053005.

35. Yip, K.M., N. Fischer, E. Paknia, A. Chari, H. Stark, 2020. Atomic-resolution protein structure determination by cryo-EM. Nature 587, 157–161.

36. Zhang, K., G.D. Pintilie, S. Li, M.F. Schmid, W. Chiu, 2020. Resolving individual atoms of protein complex by cryo-electron microscopy. Cell research 30, 1136–1139.

37. Zhu, J., Q. Zhang, H. Zhang, Z. Shi, M. Hu, C. Bao, 2023. A minority of final stacks yields superior amplitude in single-particle cryo-EM. Nature Communications 14, 7822.

